# Conserved jumbo phage factors required for protein import into a phage nucleus

**DOI:** 10.1101/2024.03.27.586873

**Authors:** Claire Kokontis, Timothy A Klein, Sukrit Silas, Joseph Bondy-Denomy

**Affiliations:** Department of Microbiology and Immunology, University of California, San Francisco, San Francisco, CA, 94158, USA; Quantitative Biosciences Institute, University of California, San Francisco, San Francisco, CA 94158, USA

## Abstract

Bacteriophages use diverse mechanisms to evade anti-phage defenses systems. ΦKZ-like jumbo phages assemble a proteinaceous nucleus-like compartment that excludes antagonistic host nucleases, while internalizing DNA replication and transcription machinery^1,2,3,4^. The phage factors required for protein import and the mechanisms of selectivity remain unknown, however. Here, we uncover an import system composed of proteins highly conserved across nucleus-forming phages, together with additional cargo-specific contributors. Using a genetic selection that forces the phage to decrease or abolish import of specific proteins, we determine that the import of five different phage nuclear-localized proteins (Nlp) all require distinct interfaces of the same factor, Imp1 (gp69). Imp1 forms discrete puncta in the phage nuclear periphery likely in complex with a direct interactor Imp6 (gp67), a conserved protein encoded nearby. The import of some proteins, including a host topoisomerase (TopA), additionally require Imp3 (gp59), a factor required for proper Imp1 function. Three additional phage proteins (Imp2, Imp4, Imp5) are also required for the import of two queried nuclear cargos, perhaps acting as specific adaptors. We therefore propose a core import system including Imp1, Imp3, and Imp6 with the highly selective Imp1 protein licensing transport through a protein lattice.

## Main

Segregation of cytoplasmic and organelle activity, with regulated movement of biomolecules between them, is a fundamental aspect of eukaryotic life. DNA-containing organelles, long thought to be exclusive to eukaryotes, have been recently discovered during bacteriophage infection in multiple genera of Gram-negative bacteria including *Pseudomonas, Escherichia, Serratia, Erwinia, Vibrio,* and *Salmonella*^1,2,4–8^. A phage-produced proteinaceous nucleus-like structure compartmentalizes DNA replication and transcription, while translation occurs in the cytoplasm^1^. This compartment is primarily made of a single protein called Chimallin or PhuN and excludes host nucleases, such as CRISPR-Cas and restriction enzymes^3^, but imports various phage proteins and at least one host protein^1,9^. How this selectivity is achieved and the mechanism of movement through the protein-based lattice^5,10,11^ that forms the phage nucleus remain unaddressed. Understanding how proteins move from the cytoplasm into the phage nucleus, analogous to what is achieved by secretion systems or eukaryotic nuclear import, may reveal new fundamental biological mechanisms.

Transport of cargo proteins through the eukaryotic lipid nuclear membrane is mediated by binding between linear sequence motifs and adaptor proteins called importins that shuttle them through a massive ∼100 MDa nuclear pore^12^. In the ΦKZ-like jumbo phages, there is no known signal that licenses proteins for import, no known adaptors, and no confirmed nuclear shell constituents that function as an entry point for phage proteins. Given the abundance of uncharacterized genes in the large ΦKZ-like genomes (i.e. > 200 kb) and the general insolubility of the protein-based nucleus limiting basic interaction approaches^13^, we use an unbiased genetic selection to identify the requirements for protein import. The screen implicates three proteins (Imp1, Imp3, Imp6) that are broadly conserved in nucleus-forming phages with a role in protein import and an additional three (Imp2, Imp4, Imp5) that appear to be conserved only among related *Pseudomonas*-infecting nucleus-forming phages. These proteins lack predicted molecular functions, and therefore form a unique phage protein trafficking system that enables transit into the phage nucleus.

### Identification of a phage protein required for nuclear cargo import

The ΦKZ phage nucleus excludes the restriction enzyme EcoRI (278 amino acids, 31.5 kDa), but fusion of EcoRI to a <u>n</u>uclear-localized <u>p</u>rotein gp152 (UvsX/RecA homolog, hereafter referred to as Nlp1) allows EcoRI to enter the nucleus and cleave phage DNA^3^. This suggests that import is licensed by recognition of nuclear cargo, rather than acting by excluding antagonistic host proteins. To identify phage factors that are required for proteins to move into the nucleus after synthesis in the cytoplasm, we used a genetic selection experiment. We individually fused four nuclear proteins, Nlp1 (gp152, RecA-like), Nlp2 (gp155, RNase H-like), Nlp3 (gp104), and Nlp4 (gp171) to the C-terminus of EcoRI-sfCherry2 constructs (hereafter referred to as EcoRI-NlpX fusions) to impart selective pressure on the phage to reduce or abolish import of these cargo via mutations in putative import factors^1,9^ (Fig. 1a). All fusions reduced phage titer by ∼10^6^-fold due to the 92 EcoRI recognition sites in the phage genome, while dead EcoRI (E111G) fused to Nlp1 also localizes inside the phage nucleus but provides no reduction in titer^3^. EcoRI fusions to other nuclear proteins (gp50, gp70, gp118, gp123, or gp179)^1,9^ were inactive (Extended Data Fig. 1a). An unmutagenized phage population with a titer of ∼10^11^ pfu/mL was subjected to targeting by each of the four EcoRI fusions and spontaneous mutant escape phages emerged at a frequency of ∼10^-6^-10^-7^. EcoRI-Nlp2, EcoRI-Nlp3, and EcoRI-Nlp4 fusions all selected for distinct mutations in a single uncharacterized phage gene, *orf69*, hereafter referred to as *imp1* for import gene 1 (Fig. 1b, Table 1, Extended Data Table 1). A total of 8 distinct missense mutant alleles in *imp1* were identified in this way, with no overlap (Table 1, Extended Data Table 1). Complementation with WT *imp1* expressed *in trans* allowed EcoRI-Nlp2/3/4 fusions to restrict *imp1* mutant escape phages again (Extended Data Fig. 1b), demonstrating that mutations in *imp1* are causal for escaping each EcoRI-Nlp fusion.

**Figure 1:**
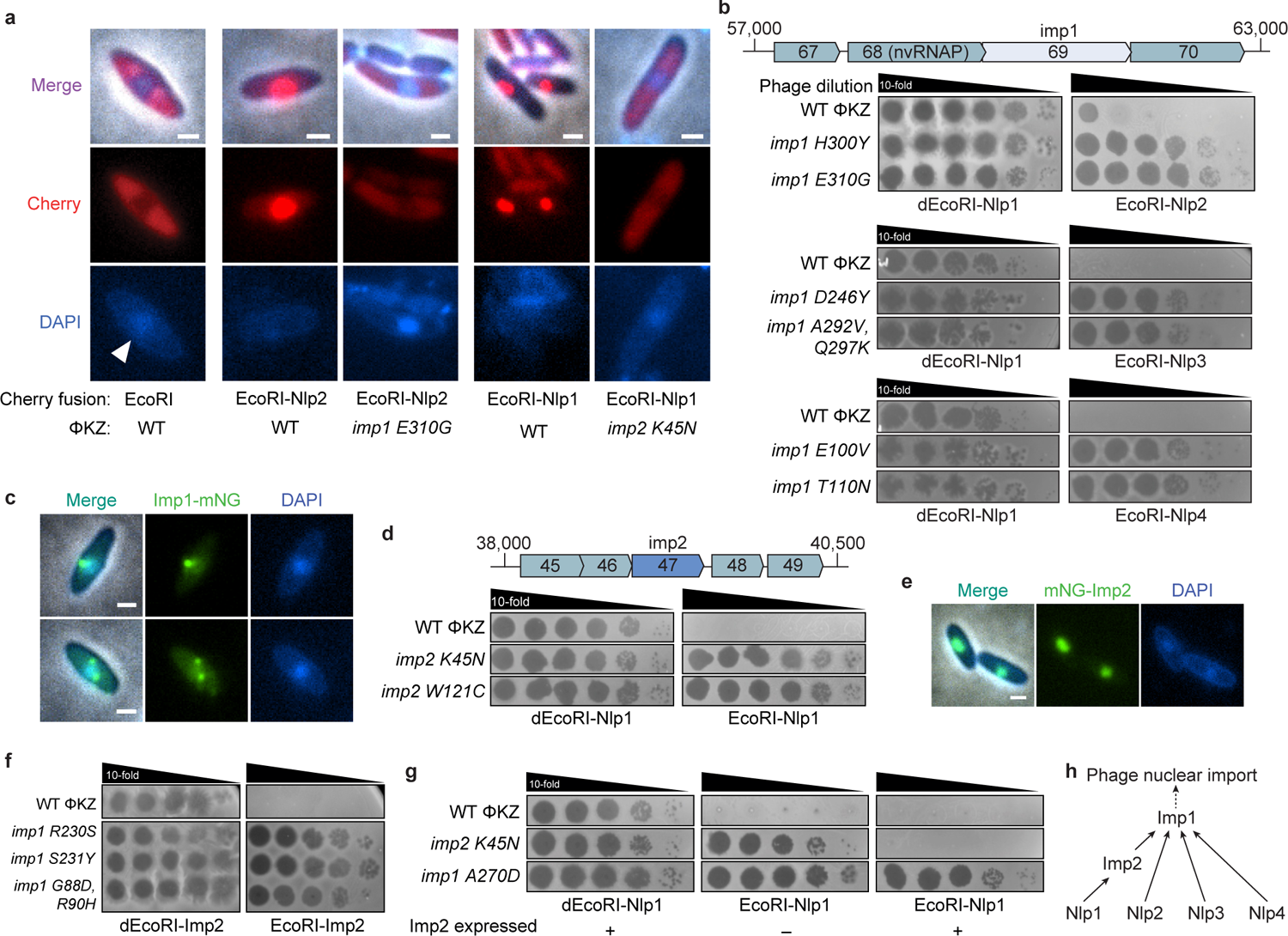
Isolation of mutant phages that reduce protein import into phage nucleus **a**, Live-cell fluorescence microscopy of *P. aeruginosa* strain PAO1 expressing the indicated sfCherry2 fusion proteins infected with WT ΦKZ or the indicated phage mutant. DAPI stain indicates phage DNA within the phage nucleus (white arrow). **b**, **d**, **f**, Plaque assays with the indicated WT or mutant phage spotted in 10-fold serial dilutions on a lawn of PAO1 expressing indicated sfCherry2 fusion. dEcoRI, catalytically inactive EcoRI. **c**, **e**, Live-cell fluorescence microscopy of Imp1 fused to mNeonGreen (mNG) infected with WT ΦKZ (n = 94) (**c**) or Imp2 fused to mNeonGreen and infected with WT ΦKZ (n = 129) (**e**). **g**, Plaque assays with the indicated WT or mutant phage spotted in 10-fold serial dilutions on a lawn of PAO1 expressing the indicated sfCherry2 fusion, with or without expression of Imp2 *in trans* (+/– Imp2). **h**, Schematic representing the factors required for nuclear import of proteins queried in the EcoRI selections, as determined by the escape mutations isolated in *imp1* and *imp2* from each selection (i.e. Nlp2 → Imp1 indicates Nlp2 requires WT Imp1 for its nuclear import). All plaque assays were replicated three or more times. All microscopy experiments were replicated two or more times. Scale bars, 1µm.

**Table 1:** Summary of mutant alleles selected for by EcoRI fusions.

| **Selection** | **Mutated gene** | **Accession no.** | **No. of unique alleles** |
| --- | --- | --- | --- |
| EcoRI-Nlp1 | *imp2 (orf47)* | NP_803613.1 | 8 |
| Tn7::Imp2<br>[EcoRI-Nlp1] | *imp1 (orf69)* | NP_803635.1 | 1 |
| EcoRI-Nlp2 | *imp1* |  | 3 |
| EcoRI-Nlp3 | *imp1* |  | 2 |
| EcoRI-Nlp4 | *imp1* |  | 3 |
| EcoRI-Imp2 | *imp1* |  | 14 |
|  | *imp3 (orf59)* | NP_803625.1 | 1 |
| EcoRI-Nlp2+<br>EcoRI-Imp2* | *imp1* |  | 10 |
| Tn7::Imp1<br>[EcoRI-Imp2] | *imp1* |  | 2 |
|  | *imp3* |  | 9 |
| EcoRI-Imp1<sub>ΦPA3</sub> | *imp1* |  | 2 |
|  | *imp3* |  | 4 |
| EcoRI-TopA | *imp1* |  | 2 |
|  | *imp3* |  | 2 |
|  | *imp4 (orf48)* | NP_803614.1 | 4 |
|  | *imp5 (orf287)* | NP_803853.1 | 1 |

\* indicates selections were performed sequentially

To confirm that mutations in *imp1* decreased protein import, we imaged EcoRI-Nlp2 during infection with WT phage or an *imp1 E310G* mutant. Indeed, protein import was substantially decreased by this mutation (Fig. 1a, Extended Data Fig. 2b-c). We next determined the subcellular localization of Imp1 with an mNeonGreen fusion expressed from the host chromosome during infection with WT ΦKZ. Upon infection, Imp1 went from being diffuse in the cell to forming 1-2 puncta at the periphery of the phage nucleus (Fig. 1c, Extended Data Fig. 3a), a localization consistent with a role in protein import. These tagged proteins, however, were not able to resensitize imp1 mutant phages to EcoRI-Imp2 targeting (Extended Data Fig. 3b). This suggests that Imp1-mNG is inactive, perhaps because the fusion blocks interactions with cargo or other binding partners. The ΦPA3 homolog of Imp1 (ΦPA3 gp63) was also recently identified as an interactor with Chimallin/PhuN through proximity labeling in phage ΦPA3^9^. Microscopy experiments with the ΦPA3 Imp1 homolog as well the phage 201Φ2-1 Imp1 homolog (201-Φ2-1 gp125) similarly revealed puncta in the nuclear periphery^1,9^. Together, these data suggest that Imp1 plays an important role in protein import into the phage nucleus.

Phages that escaped EcoRI-Nlp1 restriction, on the other hand, had 8 distinct missense or nonsense mutations in a second uncharacterized gene, *orf47* (*imp2*) (Fig. 1d, Table 1, Extended Data Table 1). EcoRI-Nlp1 import into the phage nucleus was indeed disrupted during infection with an *imp2 K45N* mutant phage (Fig. 1a, Extended Data Fig. 2d-e). *imp2* expression *in trans* also restored EcoRI-Nlp1 import and targeting of the phage, confirming causality of this mutation (Extended Data Fig. 1c). Imp2 only appears to be conserved among related *Pseudomonas*-infecting nucleus-forming phages based on sequence searches, and none of its homologs have any known function (Extended Data Fig. 1d). The importance of Imp2 in Nlp1 import was confirmed by targeting a related phage, ΦPA3, with EcoRI-Nlp1, which similarly revealed phage escape mutations in *orf44*, encoding an Imp2 homolog (Extended Data Fig. 1e). Since Imp2 mutations were uniquely selected for by EcoRI-Nlp1, we next wondered if Imp2 was also an imported protein. We fused Imp2 to mNeonGreen to assess its localization during phage infection and found that it is indeed imported into the nucleus (Fig. 1e). This suggests that Imp2 is required for Nlp1 import but is itself imported. We therefore repeated the selection, fusing EcoRI to Imp2, which resulted in robust phage targeting while dead EcoRI-Imp2 did not. We found that phages that escaped EcoRI-Imp2 targeting contained new missense mutations in *imp1* (Fig. 1f, Table 1, Extended Data Table 1). To understand why mutations in *imp1* were not initially seen under EcoRI-Nlp1 selection, we expressed Imp2 *in trans,* preventing successful phage escape via mutations in this gene. Indeed, this approach selected directly for a mutation in *imp1* (*A270D*) (Fig. 1g). This new class of Nlp1 import-deficient mutant phage emerged at a lower frequency of ∼10^-7^, explaining why it wasn’t initially seen. Together, these data demonstrate that Nlp1 uniquely requires Imp2 for its import, while import of Nlp1-Nlp4 and Imp2 all require Imp1 (Fig. 1h).

### Imp1 has distinct interfaces required for import of different phage nuclear cargo

The selective pressure to reduce import of Nlp1 and Imp2 selected for 15 unique mutant alleles in *imp1*, none overlapping with the 8 *imp1* mutant alleles previously isolated by the Nlp2, Nlp3, or Nlp4 fusions (Table 1, Extended Data Table 1). Given the convergence on Imp1, we next queried whether the unique mutations decreased import of all cargo or only some. Cells expressing each of the EcoRI-Nlp fusions were infected with WT ΦKZ or various *imp1* mutant phages. Interestingly, individual *imp1* mutations only decreased the phage targeting activity of the EcoRI fusion construct they were selected under, suggesting specific perturbations to import (Fig. 2a). This suggests that the *imp1* mutation acquired by each escape phage likely maintains sufficient expression, localization and import function for other cargo while only specifically perturbing the import of one protein. To visualize the location of functional residues, an AlphaFold2 predicted structural model of Imp1 was generated, and each set of cargo-specific mutations labeled (Fig. 2b). The four imported cargo selected for Imp1 mutations that generally clustered into five regions on the Imp1 model. Consistent with discrete functional interfaces on Imp1, double mutants could be isolated by iterative exposures to different EcoRI-Nlp fusions. For example, an Imp2 import-deficient mutant phage (*imp1 G88D/R90H*) acquired a new mutation in *imp1* (*R306L*) when selected on EcoRI-Nlp2 (Fig. 2c). The inverse was also true, a phage that resisted import of EcoRI-Nlp2 (*imp1 H300Y*) acquired a second mutation in *imp1* (*I226T*) when selected on EcoRI-Imp2. Together, we propose that Imp1 is equipped with distinct interfaces to achieve import specificity and licensing.

**Figure 2:**
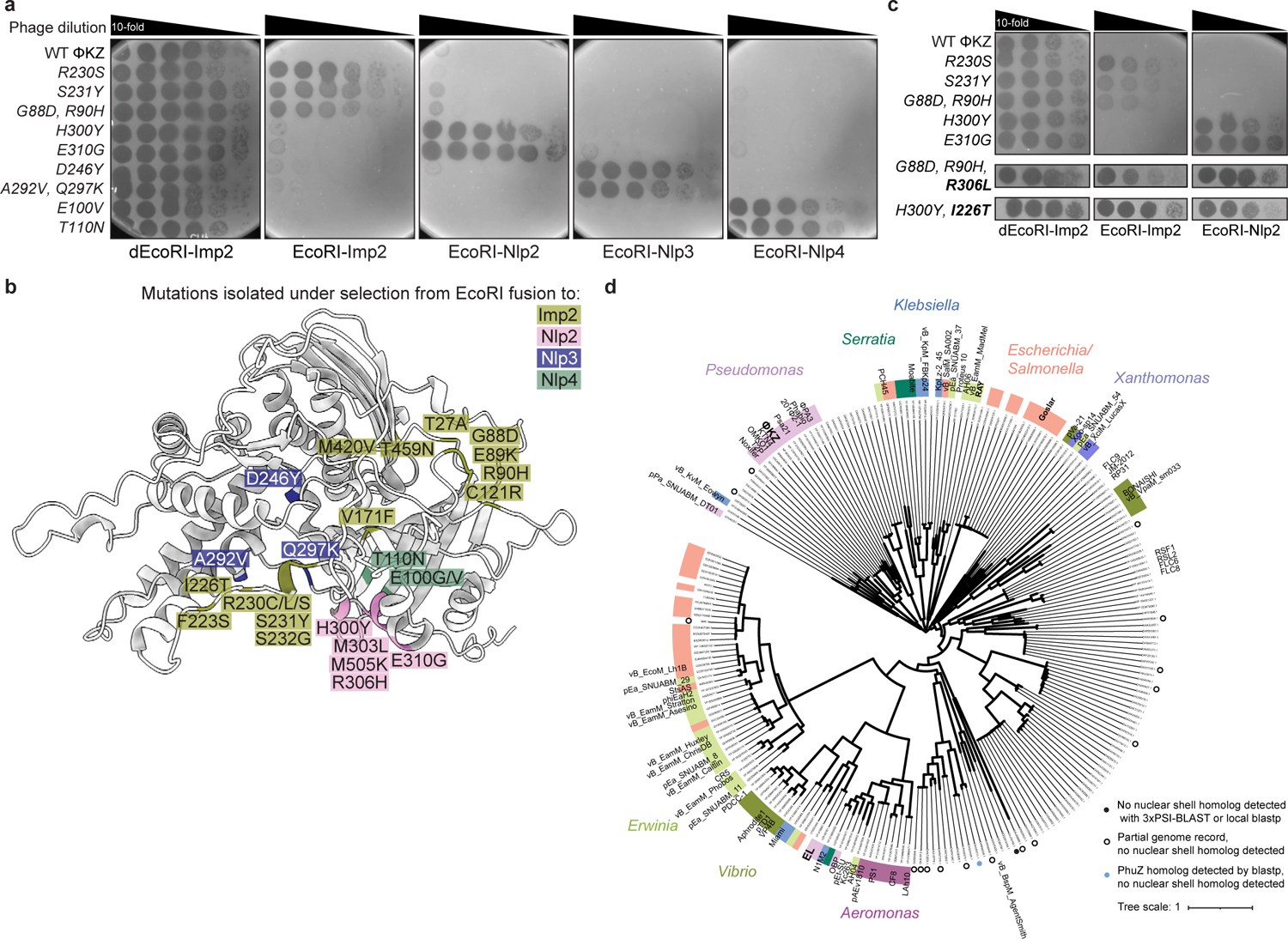
Imp1 possesses distinct interfaces to enable import specificity **a**, **c**, Plaque assays with indicated WT or *imp1* mutant phages spotted in 10-fold serial dilutions on a lawn of PAO1 expressing the indicated sfCherry2 fusion. dEcoRI, catalytically inactive EcoRI. Plaque assays were conducted as in Fig 1b. **b**, AlphaFold2 high confidence predicted structural model of Imp1, with mutated residues from isolated *imp1* mutant phages color coded by the EcoRI selection from which they were isolated. **d**, Phylogenetic tree of Imp1 homologs. A closed black circle indicates that no phage nuclear shell homolog was detected in the corresponding phage genome by 3 iterations of PSI-BLAST or local blastp. Open black circles indicate that the associated genome record is incomplete, and no phage nuclear shell homolog was detected by the same methods. A closed circle indicates that no nuclear shell homolog was detected by the same methods, but the tubulin homolog responsible for centering the phage nucleus in the cell, PhuZ, was detected by blastp. All plaque assays were replicated three or more times.

Sequence and structure-based searches with Imp1 did not reveal homologs with any known molecular function. Imp1 is widely conserved in nucleus-forming phages infecting diverse genera including *Pseudomonas, Serratia, Xanthomonas, Aeromonas, Vibrio, Erwinia,* and members of the Enterobacteriaceae family (Fig. 2d). Nearly every phage genome encoding an Imp1 protein also encodes a homolog of ΦKZ gp54/Chimallin/PhuN, the major phage nucleus protein. Only a few exceptions are noted and likely due to incomplete genomic records. Together with the genetic data demonstrating the importance of multiple Imp1 interfaces in the import of distinct cargo, its localization during infection, and its broad conservation in nucleus-forming phages, we propose that Imp1 is a key determinant of protein transport into the phage nucleus for this phage family.

### Imp1 genetic and physical interactions with *imp3* and Imp6

To identify factors required for Imp1 function or localization, we fused EcoRI (or dead EcoRI) to Imp1, to assess its proximity to phage DNA and select for mutants that disrupt its localization. Dead EcoRI-Imp1 did not reduce phage titer, while active EcoRI-Imp1 abolished phage replication such that no escape mutants could be isolated (limit of detection: 2 x 10^-9^) (Extended Data Fig. 4a). This demonstrates that the Imp1 N-terminus is localized inside the phage nucleus and suggests that mutations that alter this localization may be lethal. Fusing EcoRI to 11 different Imp1 mutant proteins, with representatives chosen from selection experiments with the various EcoRI-Nlp fusions, also maintained full phage targeting with no escape mutants isolated (Extended Data Fig. 4a). This suggests that mutant Imp1 proteins express and localize correctly, as proposed above.

To identify additional *imp1* genetic interactions not seen from the previous selections, we used two alternative selection approaches, which both led to the same new uncharacterized gene, *orf59* (*imp3*) (Fig. 3a-c). Using the EcoRI-Imp2 fusion that initially yielded *imp1* mutant phages, we repeated the selection but expressed WT Imp1 *in trans* to enable isolation of phages with mutations in other import factors. One class of mutant phages emerged that acquired spontaneous missense mutations in *imp1* (Extended Data Fig. 3b, Table 1, Extended Data Table 1), which are genetically dominant. However, most phage mutants that escaped this targeting construct acquired mutations in *imp3* (Fig. 3a). Mutations were either coding changes or 10 bp upstream of the ATG start, in the likely Shine-Dalgarno sequence (Fig 3c, Table 1, Extended Data Table 1). These data demonstrate that *imp3* is required for correct Imp1 function with respect to Imp2 import. In a complementary approach, we fused EcoRI to gp63 from phage ΦPA3, an Imp1 homolog with 52% a.a. identity (Imp1_ΦPA3_). EcoRI-Imp1_ΦPA3_ provided >100-fold weaker selection pressure against ΦKZ than EcoRI-Imp1_ΦKZ_ and enabled mutant ΦKZ phages to emerge (Fig. 3b, Table 1, Extended Data Table 1). This approach also selected for dominant *imp1* mutations and mutations in *imp3*, which possibly perturb Imp1_ΦPA3_ localization or assembly (Fig. 3c, Extended Data Fig. 3b).

**Figure 3:**
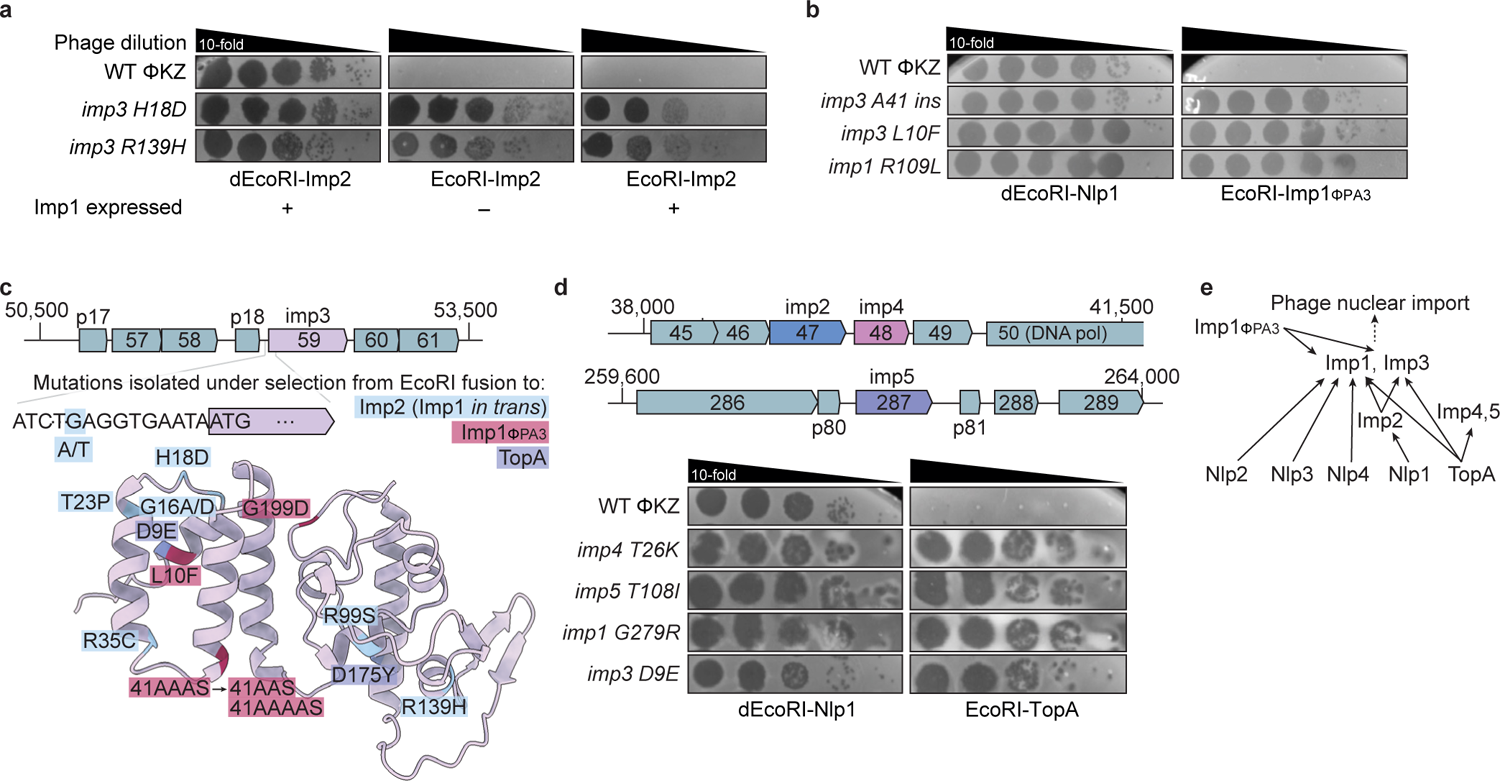
Imp3 is required for proper Imp1 function **a**, **b**, Plaque assays with the indicated WT or mutant phage spotted on PAO1 expressing the indicated sfCherry2 fusion, with or without Imp1 expressed *in trans* (+/– Imp1) (**a**); Imp1_ΦPA3_ indicates the phage ΦPA3 Imp1 homolog (**b**); dEcoRI, catalytically inactive EcoRI. Plaque assays were performed as in Fig 1b. **c,** AlphaFold2 high confidence predicted structural model of Imp3 with mutated DNA bases or residues from isolated *imp3* mutant phages color coded by the EcoRI selection from which they were isolated. **d**, Plaque assays with the indicated WT or mutant phage spotted on PAO1 expressing the indicated sfCherry2 fusion. **e**, Schematic representing the factors required for nuclear import of all proteins queried in the EcoRI selections in this work, as determined by the escape mutations isolated in *imp1*-*imp5* from each selection. All plaque assays were replicated three or more times.

Only one host protein is known to be localized inside the phage nucleus, topoisomerase TopA^1^. The EcoRI-TopA fusion also selected for escape phages (∼10^-6^ frequency) with mutations in *imp1* and *imp3* (Fig. 3c-d, Extended Data Fig. 3b). Additionally, EcoRI-TopA selected for mutations in two new uncharacterized genes, *orf48* (*imp4*) and *orf287* (*imp5*) (Fig. 3d). Like Imp2, identified above as a protein specifically required for Nlp1 import, Imp4 and Imp5 are conserved only in ΦKZ-like phages infecting *Pseudomonas* and no other EcoRI fusion selected for mutations in these genes (Extended Data Fig. 5a). Thus, we interpret this to mean that Imp4 and Imp5 are specifically required for TopA import, while core import proteins Imp1 and Imp3 are additionally required.

To understand the contribution of Imp3 to protein import, we first assessed its phylogenetic distribution and localization. Like Imp1, Imp3 is well-conserved among nucleus-forming phages beyond those that infect *Pseudomonas*^6^. However, fluorescently tagged Imp3 did not express well and thus its localization could not be assessed. Additionally, EcoRI-Imp3 fusions were inactive (data not shown) and thus whether it is proximal to phage DNA could not be ascertained. Finally, we suspect that co-encoded genes in a putative operon^14^ with *imp3* are required for proper expression or function as these genes (*p18-imp3-orf60-orf61*) were required to achieve partial complementation EcoRI-Imp1_ΦPA3_ and EcoRI-TopA escape (Extended Data Fig. 4c, 5b). The functional contributions of *imp3* and its gene neighbors will require further investigation.

Finally, we reasoned that the process of protein import may require additional proteins that associate with Imp1 that were not identified by genetic selection experiments. To find candidate interactors, we examined the operon structure and conservation of the locus around *imp1* (*orf69*). This locus encodes gp67-70 (Fig. 1b), but previous work has established that gp68 (together with gp71-73 and gp74) is part of the nvRNAP complex^15^. Therefore, we assessed whether Imp1 interacts with gp67 or gp70. 6xHis-tagged Imp1 expressed in *E. coli* immunoprecipitated with gp67-FLAG, but not gp70-FLAG (Fig. 4a) and a reciprocal FLAG immunoprecipitation revealed the same result (Fig. 4b). To confirm that Imp6-FLAG doesn’t adhere to Ni-NTA beads non-specifically, 6xHis-tagged Nlp1 was used, which did not pull-down gp67-FLAG (Fig. 4a). The Imp1-gp67 interaction was stable over size exclusion chromatography (Fig. 4c). Its elution position and mass photometry suggested a ∼120 kDa complex consistent with a stoichiometry of gp67_2_:Imp1_1_ (Fig. 4d). We therefore propose naming gp67 as Imp6, a direct binder of the core Imp1 protein and a putative member of a phage nuclear import complex.

**Figure 4:**
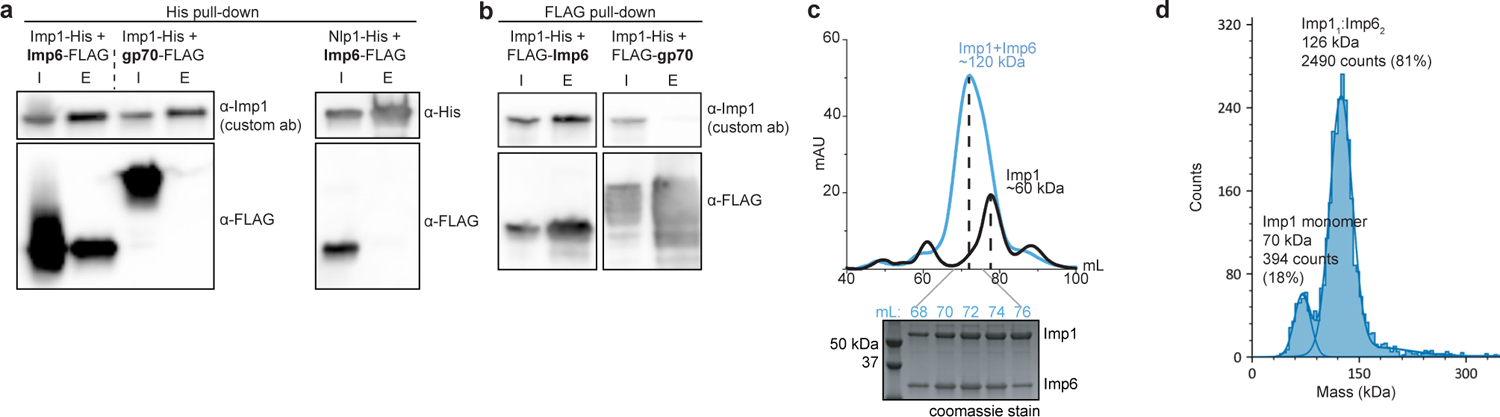
Imp6 stably binds to Imp1 **a**, **b**, Western blot analysis (anti-Imp1) of a His pull-down of His-tagged Imp1 with FLAG-tagged Imp6, gp70, or Nlp1-His with Imp6-FLAG as a negative control (**a**) or a FLAG pull-down of Imp1-His with FLAG-Imp6 or FLAG-gp70 (right) (**b**); I, input; E, elution. **c**, Size exclusion chromatography (SEC) of Imp1-His alone or in complex with Imp6-FLAG. Peak Imp1-His/Imp6-FLAG SEC fractions (indicated by mL collected off the column) were run on an SDS-PAGE gel and stained with coomassie. **d**, Estimation of Imp1-His/Imp6-FLAG complex molecular weight by mass photometry.

## Discussion

The ΦKZ-like family of jumbo phages assembles a remarkable protein-based nucleus during infections, which selectively internalizes or excludes proteins, demonstrating a eukaryotic nucleus-like segregation ability. These phages often have very large genomes (> 200 kb) with many genes of unknown function that likely contribute to this and other processes executed by this phage family, necessitating unbiased approaches to identify gene functions. Through the deployment of a genetic selection approach, we have identified 5 factors (Imp1-5) required for the import of 6 proteins (Nlp1-4, Imp2, TopA).

A key finding from this work is the functional identification of an import factor Imp1, a previously uncharacterized protein that is broadly conserved in nucleus-forming jumbo phages. Imp1 is a key specificity determinant for licensing import, as demonstrated by the isolation of individual mutations that specifically abolish the import of distinct cargo proteins. However, it currently remains unclear how proteins physically transit into the phage nucleus. Possible options include a bona fide pore, a flippase-like mechanism, or an unfoldase that participates in threading the cargo through the protein lattice. We additionally report a functional role for a protein encoded nearby, Imp3, and a direct physical interaction between Imp1 and Imp6. An Imp6 homolog in phage ΦPA3 was recently described as a non-specific RNA binding protein localized to the periphery of the phage nucleus^16^. Future work is needed to determine the connection between non-specific RNA binding and potential contributions to RNA export/localization and protein import.

The fundamental biological challenge of protein localization in segregated organellar compartments has largely been studied in eukaryotic systems. This phage family and its protein-based nucleus provide a new challenge and opportunity for understanding the basic mechanisms of protein movement. The identification of phage genes required for import, along with a method for their discovery in other phage systems, are key first steps to unraveling this fascinating mystery.

## Methods

### Bacterial growth and genetic manipulation

*E. coli* strains XL1Blue and BL21 (DE3) and *P. aeruginosa* strain PAO1 were grown in Lysogeny Broth (LB) medium at 37 °C shaking at 175 rpm. Bacteria were plated on LB solid agar and any necessary antibiotics to maintain plasmids, with 10 mM MgSO_4_ when plating for phage infection. Expression of genes inserted in the chromosomal attTn7 site was induced with 1 mM IPTG in the LB solid agar. Basal expression of EcoRI-sfCherry2-NlpX or -ImpX fusions cloned into the pHERD30T^17^ plasmid under the leaky arabinose-inducible promoter was sufficient to induce phage restriction and enable visualization by microscopy, and thus L-arabinose was not added to plates or liquid cultures.

### Phage growth

Phages were grown at 30 °C on LB solid agar with 10 mM MgSO_4_ with any necessary antibiotics and inducer. 150 µL of bacteria and 10 µL of phage were mixed in 3 mL of 0.35% top agar with 10 mM MgSO_4_ and plated on LB solid agar.

Plates were incubated at 30 °C overnight. The next day, individual plaques were picked and stored in 200 µL of SM phage buffer. Escaper phages were plaque purified three times by repeating this method. High titer lysates were generated by infecting PAO1 expressing the fusion construct that was used in selection overnight in liquid LB 10 mM MgSO_4_ with appropriate antibiotics and inducers at 37 °C. The supernatant was collected and treated with 5% volume chloroform, shaken gently for 2 min, and spun down for 5 min at max speed to remove cell debris. This was repeated once more, and the final phage lysate was stored with 1-5% volume chloroform.

### Phage spot titration plaque assays

150 µL PAO1 culture was mixed with 0.35% top agar and poured on solid LB agar plates. Once solidified, 10-fold dilutions of phage in SM phage buffer were spotted on the surface in 3 µL spots.

### Construction of fusion proteins

The shuttle vector pHERD30T^17^ was used for cloning and expression of EcoRI-sfCherry2-Nlp/Imp and mNeonGreen fusions in *P. aeruginosa* strain PAO1. The vector has a gentamicin selectable marker and an arabinose-inducible promoter. The vector was digested with NcoI and HindIII restriction enzymes and purified. Inserts were amplified by PCR from diluted ΦKZ lysates or PAO1 colonies as the DNA template, and joined into the linearized vector by Hi-Fi Gibson DNA Assembly (NEB) according to the manufacturer’s protocol. The resulting reactions were used to transform *E. coli* XL1-Blue competent cells. All plasmid constructs were sequenced using primers that annealed to the vector outside of the multiple cloning site (Quintara), or were whole-plasmid sequenced (Primordium). Plasmids were then electroporated into competent PAO1 cells and selected on gentamicin.

### Construction of chromosomal insertion strains

For chromosomal insertion of phage import factor genes into PAO1, genes of interest were cloned into the pUC18-miniTn7T-LAC^18^, once linearized with SpeI and SacI and purified. Inserts were amplified by PCR from diluted ΦKZ lysates and joined into the linearized vector by Hi-Fi DNA Assembly (NEB). The resulting vectors were used to transform *E. coli* XL1-Blues and verified by sequencing using primers that anneal to the vector outside of the multiple cloning site. The transposase helper vector pTNS3^19^ was used with miniTn7 constructs to transform PAO1 and insert the genes of interest into the PAO1 chromosome at the Tn7 locus, with a –pTNS3 control in parallel, and the transformation was selected on gentamicin. Candidate integrants were screened by colony PCR using PTn7R and PglmS-down, as well as an internal gene-specific primer paired with PglmS-up. Integrants were then amplified and sequenced with primers that annealed outside the attTn7 site to verify integration. Electrocompetent cell preparation, electroporation, integration, FLP-recombinase-mediated gentamicin marker excision using pFLP2 and plasmid curing via sucrose counterselection were performed as described previously^18^.

### Whole genome sequencing

Genomic DNA extraction from phage lysates was performed by adding 200 µL of lysis buffer (final concentration 10 mM Tris, 10 mM EDTA, 100 µg/mL Proteinase K, 100 µg/mL RNaseA, 0.5% SDS) to 200 µL high titer phage lysate (>10^9^ PFU/mL) and incubated at 37 °C for 30 min, and then 55° C for 30min. Preps were then purified by a phenol chloroform extraction followed by a chloroform extraction and then ethanol precipitated, or by using the DNA Clean & Concentrator Kit (Zymo Research). DNA was then quantified using the Qubit 4.0 Fluorometer (Life Technologies). 50-100 ng genomic DNA was used to prepare WGS libraries using the Illumina DNA Prep Kit (previously, Nextera Flex Library Prep Kit). A modified protocol was used, with 5x reduced quantities of tagmentation reagents per prep, except for the bead washing step where the recommended 100 µL of Tagment Wash Buffer (TWB) was used. On-bead PCR indexing-amplification was performed using custom-ordered indexing primers (IDT) matching the Illumina Nextera Index Kit sequences and 2x Phusion Master Mix (NEB). PCR reactions were amplified for 9-11 cycles, and subsequently resolved by agarose gel electrophoresis. DNA products were excised around the 400 bp size range and purified using the Zymo Gel DNA Recovery kit (Zymo Research). Libraries were quantified by Qubit. Libraries were pooled in equimolar ratios and sequenced on the Illumina MiSeq using 150-cycle v3 reagents (Single end: 150 cycles, Read 1; 8 cycles, Index 1; 8 cycles, Index 2.

Paired end: 75 cycles, Read 1; 8 cycles, Index 1; 8 cycles, Index 2, 75 cycles, Read 2). Data were demultiplexed on-instrument, and trimmed using cutadapt (v1.15) to remove Nextera adapters. Trimmed reads were mapped using Bowtie 2.0^20^. (–very-sensitive-local alignments) and alignments were visualized using IGV (v2.11.0). Mutations were called if they were present in >90% sequencing reads at loci with at least 20x coverage.

### Live-cell fluorescence microscopy

0.8% LB agar pads (25% LB, 2.5 mM MgSO_4_) were supplemented with 0.5 μg/mL DAPI for phage DNA staining. PAO1 strains expressing each of the fluorescent protein constructs were grown in liquid culture supplemented with inducer if necessary (0.05% arabinose for pHERD30T-mNG-Imp2, 0.1 mM IPTG for Tn7::Imp1-mNG) to induce construct expression until OD600 ∼0.5, and subsequently infected with ΦKZ lysate for 50 min at 30 °C before imaging. Microscopy was performed on an inverted epifluorescence (Ti2-E, Nikon, Tokyo, Japan) with the Perfect Focus System (PFS) and a Photometrics Prime 95B 25 mm camera. Images were acquired using Nikon Elements AR software (version 5.02.00). Cells were imaged through channels of blue (DAPI, 50 ms exposure, for phage DNA), green (FITC, 200 ms exposure, for mNeonGreen constructs), red (Cherry, 200 ms exposure, for sfCherry2 constructs) and phase contrast (200 ms exposure, for cell recognition) at 100x objective magnification. Final figure images were prepared in Fiji (version 2.1.0/1.53c)^21^.

### Phylogenetic analysis

Homologs of ΦKZ import factors were identified by 3 iterations of PSI-BLAST against the non-redundant protein database. Hits with >70% coverage and E-value <0.005 were included to generate a multiple sequence alignment (MSA) using MAFFT (v7.490, fast strategy). The genomes not labeled as bacteriophages with homologs were manually inspected for the presence of phage structural genes (i.e. to exclude bacterial contigs), and were excluded from the MSA input if no phage genes were annotated. After manual inspection of the MSA, results were input to FastTree (v2.1.11 SSE3, default settings) to generate a phylogenetic tree and visualized in iTOL: Interactive Tree of Life. To determine whether phage genomes containing Imp1 homologs also encode Chimallin/PhuN (gp54 in ΦKZ), gp54 homologs were acquired by 3 iterations of PSI-BLAST as described above, and genomes from Imp1 homologs were inspected for the presence of gp54 homologs. Genomes lacking an apparent gp54 homolog according to PSI-BLAST were inspected manually to identify a predicted gp54 locus (flanked by two well-conserved phage-encoded genes: DNA polymerase upstream and RNA polymerase beta subunit downstream). These candidates were confirmed as gp54 homologs by their similarity (>50% identity, >70% coverage) to a homolog present in a genome on the Imp1 tree.

### Protein expression and purification

6xHis-tagged Imp1 and Nlp1 were cloned into pET29b while FLAG-tagged Imp6 and gp70 were cloned into pETduet-1. Binary interactions were assessed by co-expression in BL21 (DE3) cells and purification by affinity chromatography using Ni-NTA. For expression, 100 mL cultures of cells were grown in LB supplemented with Kanamycin (50 µg/mL) and Ampicillin (100 µg/mL) at 37 °C shaking at 175 rpm. When the cultures reached OD600 ∼0.6, protein expression was induced with 1 mM IPTG and incubated at 18 °C for ∼16 hours. Cells were centrifuged at 5,000 x g and resuspended in lysis buffer (25 mM Tris-HCl pH 8.0, 300 mM NaCl, 10 mM imidazole, 0.5 mM TCEP, protease inhibitor). Lysis was performed by sonication at 20% amplitude for 10 s, four times. Insoluble material was removed from the lysate by centrifugation at 21,000 x g for 30 min. 300 µL of Ni-NTA resin slurry (Qiagen) was washed with 10 mL of wash buffer (25 mM Tris-HCl pH 8.0, 300 mM NaCl, 10 mM imidazole, 0.5 mM TCEP) in a gravity column. Cleared lysate was then run over Ni-NTA resin and non-specific interactors were removed by 10 mL washes using wash buffer (four washes total). Protein was eluted with 300 µL of elution buffer (25 mM Tris-HCl pH 8.0, 300 mM NaCl, 400 mM imidazole). Pulldown experiments on FLAG-tagged proteins were done similarly to those of His-tagged proteins but with several modifications. Cells were resuspended in lysis/wash buffer (25 mM Tris-HCl, pH 8.0, 300 mM NaCl, protease inhibitor) and sonicated as described above. 25 µL of anti-FLAG magnetic agarose resin (Pierce) was washed and then incubated with cleared lysate for one hour at 4 °C with constant agitation. Protein bound resin was washed five times with 1 mL of wash buffer using a magnetic separation rack. Bound protein was eluted with 100 µL of 1.5 mg/mL FLAG peptide (Millipore) resuspended in wash buffer. Protein interactions were assessed by Western blot (see below). Size exclusion chromatography was performed with a Superdex 200 increase 10/300 GL column using an AKTA Pure Protein Purification System (Cytiva). SEC buffer (25 mM Tris-HCl pH 8.0, 300 mM NaCl) was used for purification and protein samples collected were subject to Western blotting and mass photometry.

### Western blotting

The protein samples collected through pulldown and SEC analyses were mixed 3:1 with 4x Laemmli buffer supplemented with β-mercaptoethanol and boiled for 10 minutes. These samples were run on precast SDS-PAGE gels (Bio-Rad) and transferred to PVDF membranes. The membranes were blocked with TBS-T buffer (1x TBS, 0.1% Tween-20) supplemented with 5% skim milk. A custom primary antibody for Imp1 (α-Imp1, Genscript), a commercial primary FLAG antibody (α-FLAG, Millipore cat# F1804), or a commercial primary His antibody (α-His, Cell Signaling cat# 2365S) were then added to the skim milk buffer at a titer of 1:5000 and left to incubate at room temperature for one hour. The membranes were washed 3x with 10 mL of TBS-T, then incubated with α-Rabbit (Cell Signaling, cat# 7074S) or α-Mouse (Invitrogen, cat# 62-6520) secondary antibody at a titer of 1:5000 for 45 minutes at room temperature. The membrane was washed 3x with 10 mL of TBS-T and developed with Clarity Max ECL substrate (Bio-Rad). Western blot images were taken with an Azure Biosystems C400 imager.

### Mass photometry

Imp1-Imp6 protein complexes purified by size exclusion chromatography were analyzed by using a OneMP mass photometer (Refeyn). Adequate data collection was made by mixing 1 µL of 1 µM protein with 15 µL of buffer (25 mM Tris-HCl, 300 mM NaCl). Data collection was done with the AcquireMP software (Refeyn 2024 R1.1). Data was collected for one minute and yielded 3055 measurable events. Data processing was done using the DiscoverMP software (Refeyn DiscoverMP 2024 R1).

### Materials and correspondence

Supplementary Information is available for this paper. Correspondence and requests for materials should be addressed to Joseph Bondy-Denomy.

## Acknowledgments

The authors thank Bondy-Denomy lab members for input into this work, including Iana Fedorova for help running FastTree. We additionally thank David Agard, Eliza Nieweglowska, Carol Gross, and Alan Davidson for vital input.

## Author contributions

C.K. and J.B.-D. conceived the project and designed experiments. C.K. executed genetic screens, sequencing, genetics, microscopy, and associated data analysis. T.K. executed biochemical experiments. S.S. provided sequencing execution and analysis. J.B.-D. and C.K. wrote the manuscript and all authors edited. J.B.-D. supervised experiments and procured funding.

## Declaration of interests

J.B.-D. is a scientific advisory board member of SNIPR Biome and Excision Biotherapeutics, a consultant to LeapFrog Bio, and a scientific advisory board member and co-founder of Acrigen Biosciences. The Bondy-Denomy lab received research support from Felix Biotechnology.

**Extended Data Fig. 1 a-c**, Plaque assays with WT ΦKZ or WT 14-1, an EcoRI-sensitive phage (**a**) or indicated WT ΦKZ and mutant phage (**b, c**) spotted in 10-fold serial dilutions on a lawn of PAO1 expressing indicated sfCherry2 fusions or an empty vector (EV) control, with or without expression of the appropriate phage gene *in trans* from the bacterial attTn7 site (Tn7::*impX*) (**b, c**). **d**, Imp2 phylogenetic tree. **e**, Plaque assays with the indicated ΦPA3 WT or mutant phage spotted in 10-fold serial dilutions on a lawn of PAO1 expressing indicated sfCherry2 fusions. Plaque assays were performed as in Fig 1b and replicated two times or more.

**Extended Data Fig. 2 a-e**, Representative images of live-cell fluorescence microscopy of PAO1 expressing the indicated sfCherry2 fusions, infected with WT or indicated mutant ΦKZ. Scale bars, 1 µm. Microscopy was performed as in Fig. 1c and replicated two times or more.

**Extended Data Fig. 3 a**, Live-cell fluorescence microscopy of PAO1 expressing Imp1-mNeonGreen (Imp1-mNG) from the attTn7 site, infected with WT ΦKZ (left, middle column; scale bar, 1 µm) or uninfected (right column; scale bar, 2 µm). Microscopy was performed as in Fig 1c, and replicated three times. **b**, Plaque assays with the indicated WT or mutant phage spotted in 10-fold serial dilutions on a lawn of PAO1 expressing the indicated sfCherry2 fusions, with or without expression of the appropriate phage gene *in trans* from the bacterial attTn7 site (Tn7::ImpX). Plaque assays were performed as in Fig 1b and replicated two times or more.

**Extended Data Fig. 4 a**, Plaque assays with WT ΦKZ or WT 14-1 (EcoRI-sensitive phage) spotted in 10-fold serial dilutions on a lawn of PAO1 expressing the indicated sfCherry2 fusions. **b**, High confidence AlphaFold2 predicted structural model with mutated residues from isolated *imp1* mutant phages color coded by the EcoRI selection from which they were isolated. **c**, Plaque assays with the indicated WT or mutant phage spotted on PAO1 expressing the indicated sfCherry2 fusions, with or without Imp3 or the Imp3 operon (*p18-imp3-orf60-orf61*) expressed *in trans*. Plaque assays were performed as in Fig 1b and replicated two times or more.

**Extended Data Fig. 5 a**, Imp4 and Imp5 phylogenetic trees. **b**, Plaque assays with the indicated WT or mutant phage spotted in 10-fold serial dilutions on a lawn of PAO1 expressing the indicated sfCherry2 fusions, with or without Imp3 or the Imp3 operon (*p18-imp3-orf60-orf61*) expressed *in trans*. Plaque assays were performed as in Fig 1b and replicated three times.

## References

1. Chaikeeratisak, V. et al. Assembly of a nucleus-like structure during viral replication in bacteria. Science 355, 194–197 (2017).

2. Chaikeeratisak, V. et al. The Phage Nucleus and Tubulin Spindle Are Conserved among Large Pseudomonas Phages. Cell Rep. 20, 1563–1571 (2017).

3. Mendoza, S. D. et al. A bacteriophage nucleus-like compartment shields DNA from CRISPR nucleases. Nature 577, 244–248 (2020).

4. Malone, L. M. et al. A jumbo phage that forms a nucleus-like structure evades CRISPR--Cas DNA targeting but is vulnerable to type III RNA-based immunity. Nature microbiology 5, 48–55 (2020).

5. Laughlin, T. G. et al. Architecture and self-assembly of the jumbo bacteriophage nuclear shell. Nature 608, 429–435 (2022).

6. Prichard, A. et al. Identifying the core genome of the nucleus-forming bacteriophage family and characterization of Erwinia phage RAY. Cell Rep. 42, 112432 (2023).

7. Weintraub, S. T. et al. Global Proteomic Profiling of Salmonella Infection by a Giant Phage. J. Virol. 93, (2019).

8. Jacquemot, L. et al. Therapeutic Potential of a New Jumbo Phage That Infects Vibrio coralliilyticus, a Widespread Coral Pathogen. Front. Microbiol. 9, 2501 (2018).

9. Enustun, E. et al. Identification of the bacteriophage nucleus protein interaction network. Nat. Struct. Mol. Biol. 30, 1653–1662 (2023).

10. Nieweglowska, E. S. et al. The ΦPA3 phage nucleus is enclosed by a self-assembling 2D crystalline lattice. Nat. Commun. 14, 927 (2023).

11. Heymann, J. B. et al. The Mottled Capsid of the Salmonella Giant Phage SPN3US, a Likely Maturation Intermediate with a Novel Internal Shell. Viruses 12, (2020).

12. Lin, D. H. & Hoelz, A. The Structure of the Nuclear Pore Complex (An Update). Annu. Rev. Biochem. 88, 725–783 (2019).

13. Fossati, A. et al. Next-generation proteomics for quantitative Jumbophage-bacteria interaction mapping. Nat. Commun. 14, 5156 (2023).

14. Putzeys, L. et al. Refining the transcriptional landscapes for distinct clades of virulent phages infecting pseudomonas aeruginosa. *bioRxiv* 2023.10.13.562163 (2023) doi:10.1101/2023.10.13.562163.

15. Yakunina, M. et al. A non-canonical multisubunit RNA polymerase encoded by a giant bacteriophage. Nucleic Acids Res. 43, 10411–10420 (2015).

16. Enustun, E., et al. A phage nucleus-associated RNA-binding protein is required for jumbo phage infection. *bioRxiv* (2023) doi:10.1101/2023.09.22.559000.

17. Qiu, D., Damron, F. H., Mima, T., Schweizer, H. P. & Yu, H. D. PBAD-based shuttle vectors for functional analysis of toxic and highly regulated genes in Pseudomonas and Burkholderia spp. and other bacteria. Appl. Environ. Microbiol. 74, 7422–7426 (2008).

18. Choi, K.-H. & Schweizer, H. P. mini-Tn7 insertion in bacteria with single attTn7 sites: example Pseudomonas aeruginosa. Nat. Protoc. 1, 153–161 (2006).

19. Choi, K.-H. et al. Genetic tools for select-agent-compliant manipulation of Burkholderia pseudomallei. Appl. Environ. Microbiol. 74, 1064–1075 (2008).

20. Langmead, B. & Salzberg, S. L. Fast gapped-read alignment with Bowtie 2. Nat. Methods 9, 357–359 (2012).

21. Schindelin, J., et al. Fiji: an open-source platform for biological-image analysis. Nat. Methods 9, 676–682 (2012).

