## Supplementary figures and images for "Conserved jumbo phage factors required for protein import into a phage nucleus"

### Extended Data Figure 1

#### Extended Data Figure 1

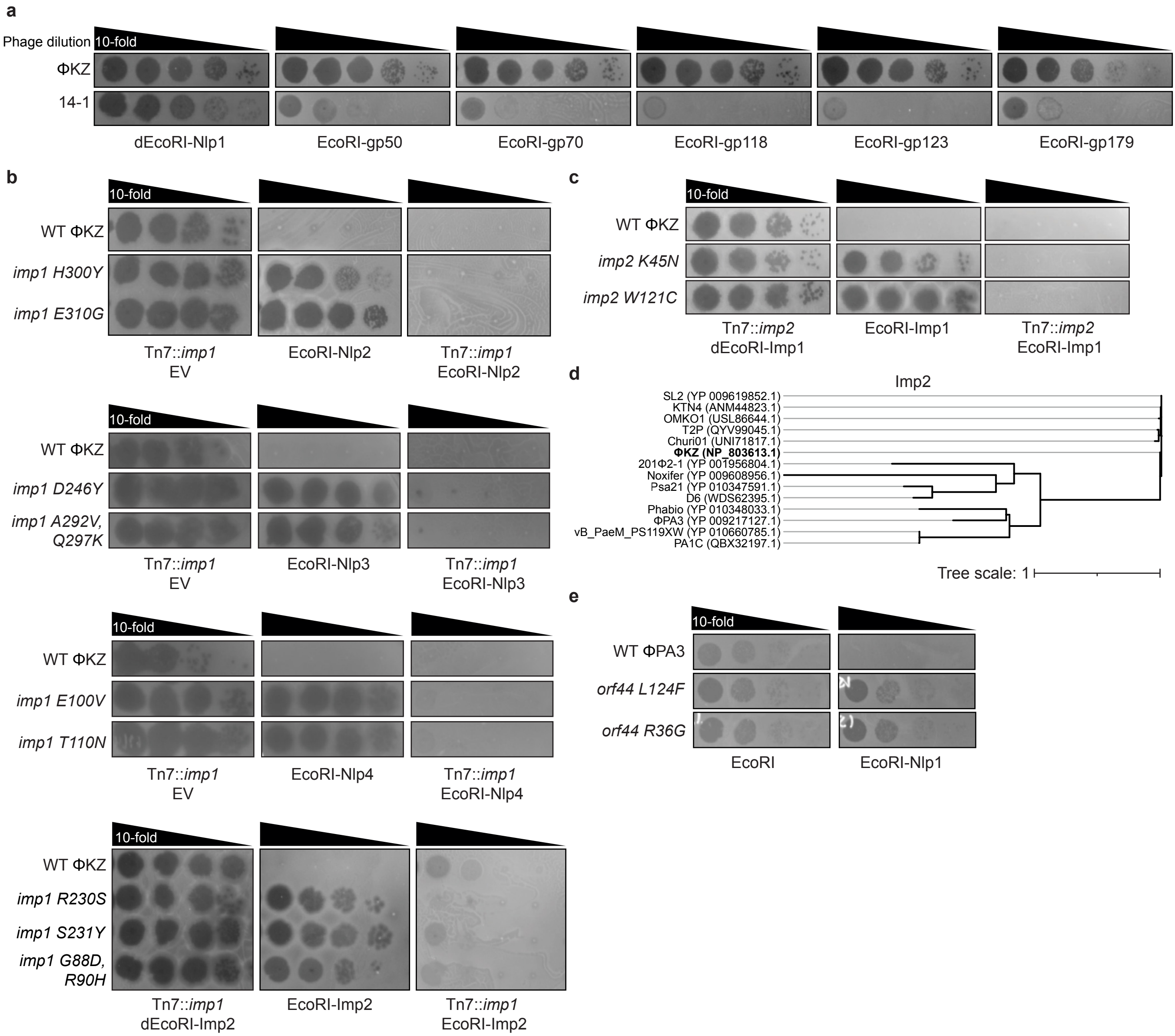

### Extended Data Figure 2

**Extended Data Figure 2**

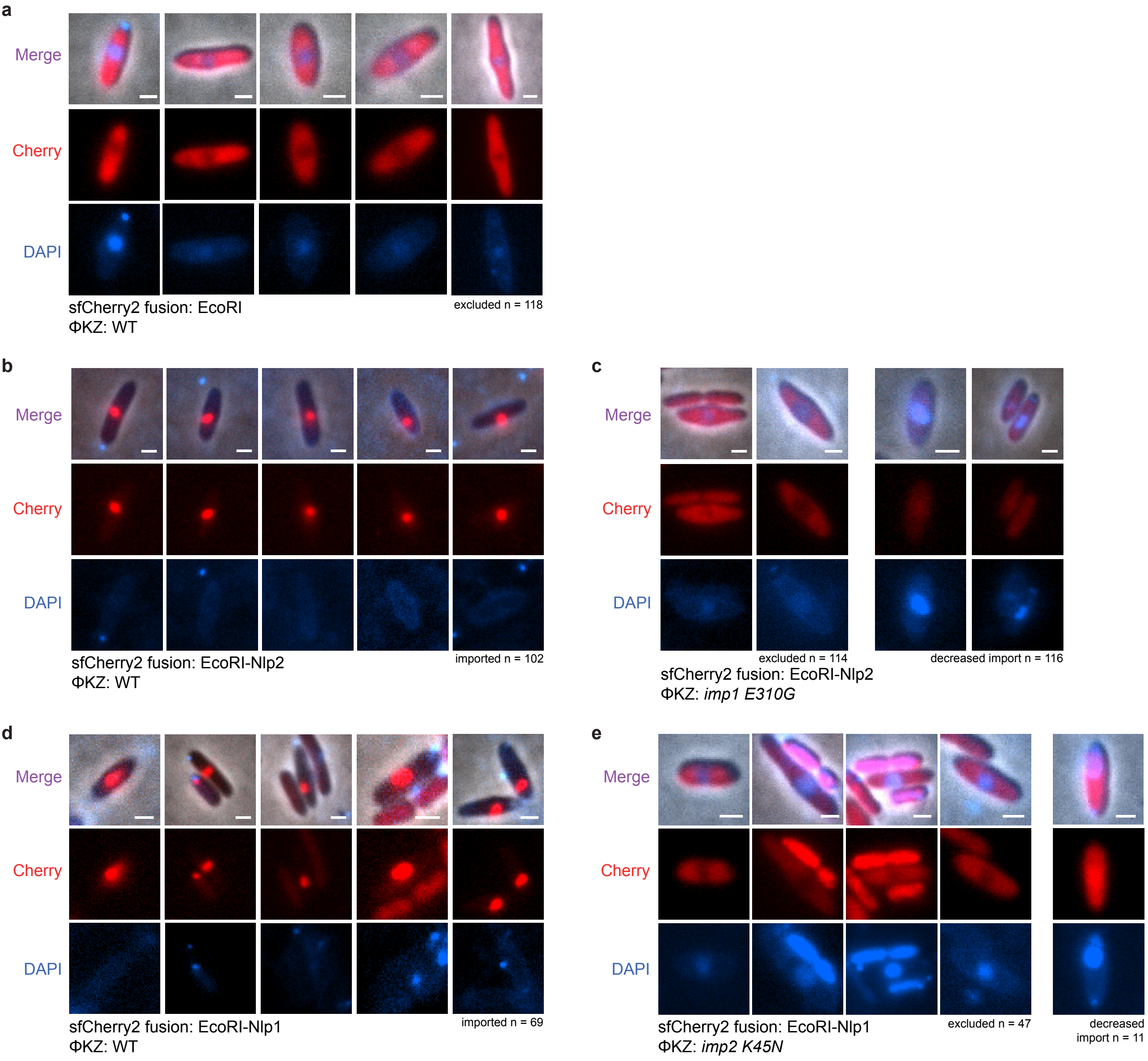

### Extended Data Figure 3

**Extended Data Figure 3**

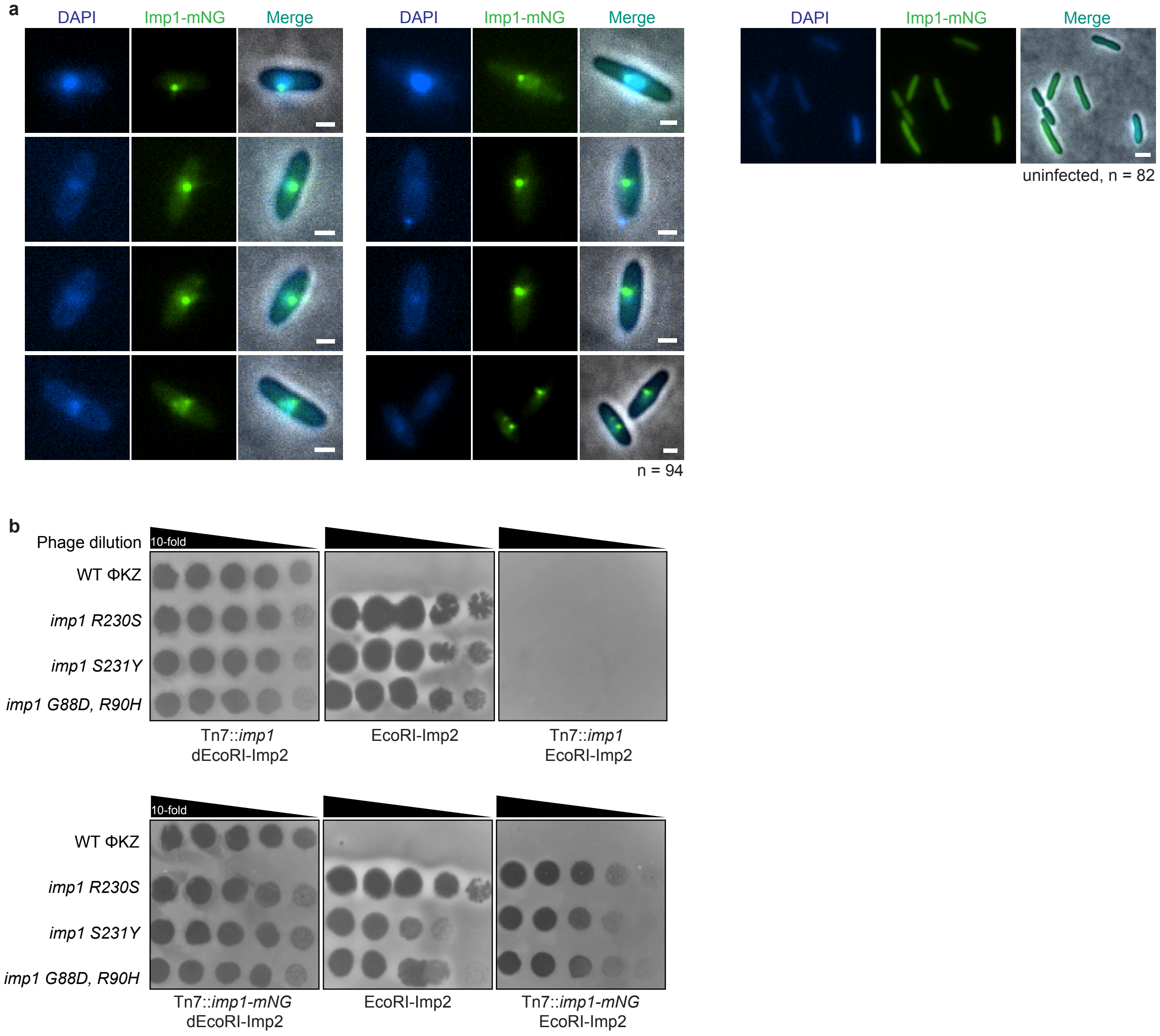

### Extended Data Figure 4

**Extended Data Figure 4**

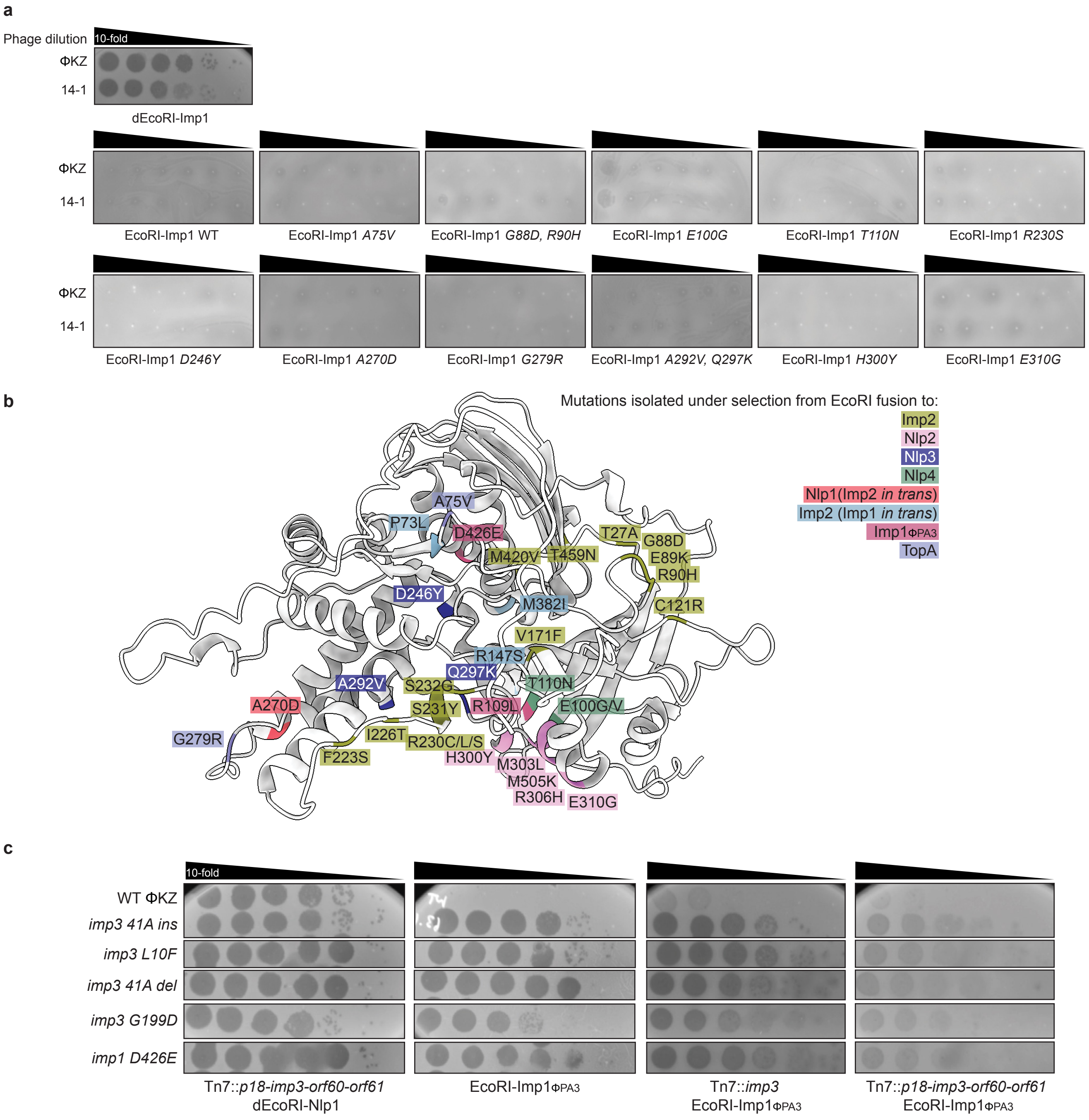

### Extended Data Figure 5

#### Extended Data Figure 5

**a**

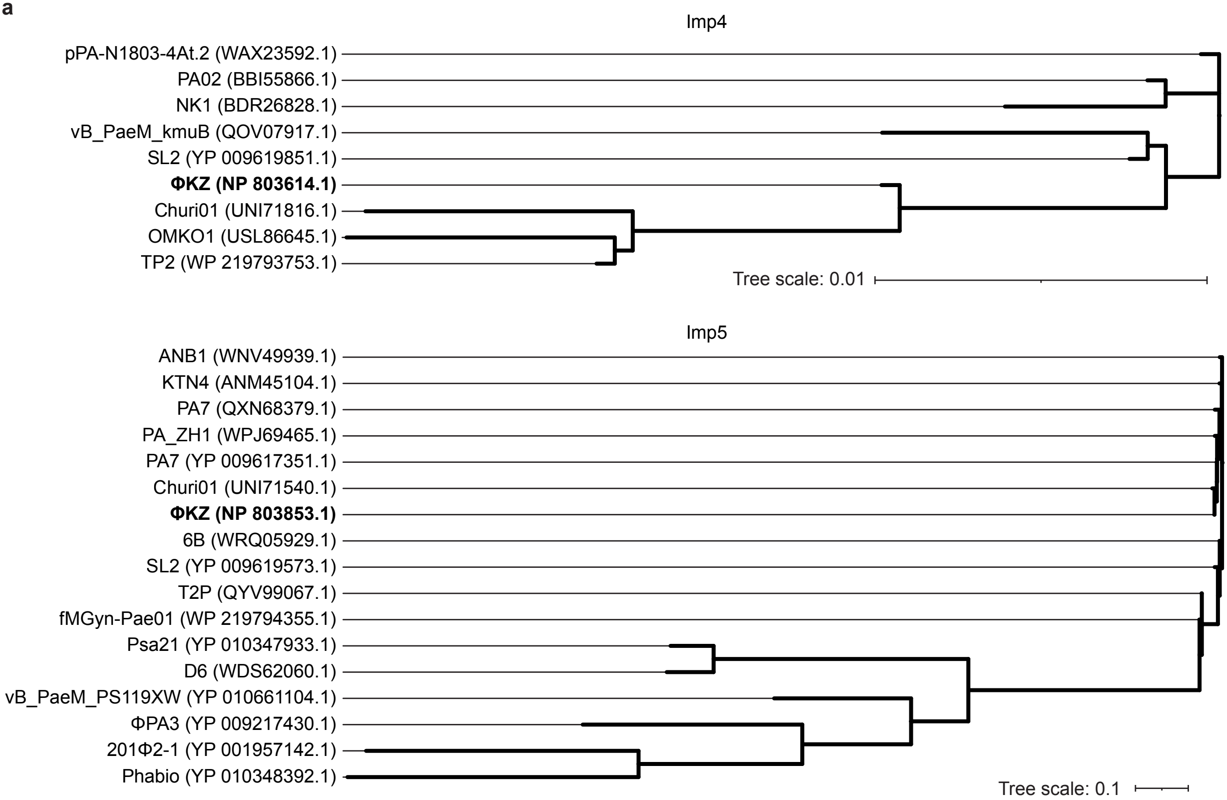


**b**

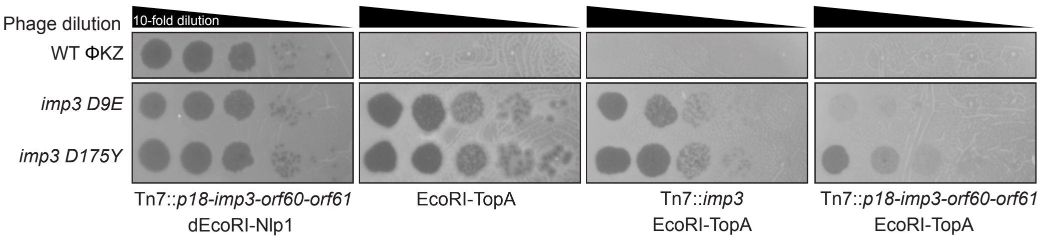
