## Extended Data Table 1 for "Conserved jumbo phage factors required for protein import into a phage nucleus"

| Selection | Escaper phage # | Mutation type | Position | Base | Codon | Gene and aa change | Sequencing method and notes |
|---|---|---|---|---|---|---|---|
| | 1 | SNP | 38578 | T-G (100%, 44 count) | cta-cga | orf46 L26R | WGS (whole genome sequencing of lysate stock) |
| | | Deletion | 38581-38934 (length 353 bp) | | | in-frame fusion of gp46-gp47 | |
| | 2 | SNP | 39338 | G-A (99%, 140 count) | gaa-aaa | orf47 E136K | WGS |
| | 3 | SNP | 39295 | G-T (100%, 110 count) | tgg-tgt | orf47 W121C | WGS |
| | 4 | SNP | 38972 | C-T (100%, 128 count) | cgg-tgg | orf47 R14W | WGS |
| | | Deletion | 52124-52126 | CTG | ctg | deletion orf59 A41 | |
| | 5 | SNP | 39005 | C-T | caa-taa | orf47 Q25STOP | PCR from lysate |
| | 6 | SNP | 39020 | A-C (100%, 118 count) | agt-cgt | orf47 S30R | WGS |
| | 7 | SNP | 38943 | C-T (100%, 23 count) | cca-cta | orf47 P4L | WGS |
| | | Insertion | 52123-52124 | CTG | ctg | orf59 insertion A41 | |
| | | Deletion | 271894-272924 | | | deletion t04-t05 | |
| | 8 | SNP | 39067 | A-C (100%, 138 count) | aaa-aac | orf47 K45N | WGS |
| EcoRI-Nlp1 | | SNP | 246718 | T-G (100%, 76 count) | noncoding region between orf259 and orf260 | | |
| | 1 | SNP | 52162 | T-C (99%, 76 count) | tac-cac | orf59 Y53H | WGS |
| Tn7::Imp2, [EcoRI-Nlp1] | | SNP | 60415 | C-A (100%, 124 count) | gct-gat | orf69 A270D | Note: The initial escaper plaque and 3x purified phage only contained orf69 A270D, not orf59 Y53H. orf59 Y53H arose during lysate propagation of the 3x purified plaque |
| | 1 | SNP | 60504 | C-T (100%, 79 count) | cat-tat | orf69 H300Y | WGS |
| | 2 | SNP | 60535 | A-G (100%, 184 count) | gaa-gga | orf69 E310G | WGS |
| | 3 | SNP | 60513 | A-T | atg-ttg | orf69 M303L | PCR 2x purified plaque |
| | | SNP | 60520 | T-A | atg-aag | orf69 M305K | |
| EcoRI-Nlp2 | | SNP | 60523 | G-A | cgc-cac | orf69 R306H | |
| | 1 | SNP | 60342 | G-T (100%, 257 count) | gat-tat | orf69 D246Y | WGS |
| | 2 | SNP | 60481 | C-T (91%, 148 count) | gct-gtt | orf69 A292V | WGS |
| EcoRI-Nlp3 | | SNP | 60495 | C-A (99%, 144 count) | caa-aaa | orf69 Q297K | |
| | 1 | SNP | 59935 | C-A (100%, 298 count) | act-aat | orf69 T110N | WGS |
| | 2 | SNP | 59905 | A-G (98%, 122 count) | gag-ggg | orf69 E100G | WGS |
| | | SNP | 65610 | A-G (99%, 193 count) | noncoding region between p21 and p22 | | |
| EcoRI-Nlp4 | 3 | SNP | 59905 | A-T (100%, 208 count) | gag-gtg | orf69 E100V | WGS |
| | 1 | SNP | 18020 | G-A (100%, 78 count) | gaa-aaa | orf27 F407K | WGS |
| | | SNP | 60294 | C-A (100%, 49 count) | cgc-agc | orf69 R230S | |
| | 2 | SNP | 60298 | C-A (99%, 112 count) | tcc-tac | orf69 S231Y | WGS |
| | 3 | SNP | 44131 | T-C (99%, 99 count) | ata-atg | p16 I238M | WGS |
| | | SNP | 54062 | G-A (100%, 68 count) | gct-act | orf63 A35T | |
| | | SNP | 57422 | T-C (100%, 106 count) | act-acc | orf67 T88T | |
| | | SNP | 59869 | G-A (100%, 133 count) | ggt-gat | orf69 G88D | |
| | | SNP | 59875 | G-A (100%, 128 count) | cgc-cac | orf69 R90H | |
| | | SNP | 60653 | C-T (100%, 112 count) | cgc-cgt | orf69 R349R | |
| | | SNP | 64760 | G-A (100%, 107 count) | tgg-tag | orf72 W92STOP | |
| | | SNP | 91137 | C-A (100%, 93 count) | cca-caa | p30 P9Q | |
| | | SNP | 129971 | T-C (100%, 79 count) | atc-gtc | orf128 I120V | |
| | 4 | SNP | 59685 | A-G | act-gct | orf69 T27A | PCR from lysate |
| | 5 | SNP | 59967 | T-C (100%, 99 count) | tgc-cgc | orf69 C121R | WGS |
| | 6 | SNP | 59871 | G-A | gag-aag | orf69 E89K | PCR from lysate |
| | 7 | SNP | 52057 | C-A (96%, 24 count) | cat-aat | orf59 H18N | WGS |
| | | Deletion | 70301 | T | noncoding region between orf75 and orf76 | | |
| | 8 | SNP | 60864 | A-G | atg-gtg | orf69 M420V | PCR unpurifed plaque |
| | 9 | SNP | 60117 | G-T | gtt-ttt | orf69 V171F | PCR unpurifed plaque |
| | 10 | SNP | 60283 | T-C | atc-acc | orf69 I226T | PCR unpurifed plaque |
| | 11 | SNP | 60274 | T-C | ttt-tct | orf69 F223S | PCR unpurifed plaque |
| | 12 | SNP | 60294 | C-T | cgc-tgc | orf69 R230C | PCR unpurifed plaque |

| | | | | | | | |
|---|---|---|---|---|---|---|---|
| | 13 | SNP | 60295 | G-T | cgc-ctc | orf69 R230L | PCR unpurifed plaque |
| | 14 | SNP | 60300 | A-G | agt-ggt | orf69 S232G | PCR unpurifed plaque |
| EcoRI-Imp2 | 15 | SNP | 60982 | C-A | act-aat | orf69 T459N | PCR unpurifed plaque |
| | 1 | SNP | 60301 | G-A | agt-aat | new orf69 S232N, still has H300Y | PCR unpurified plaque |
| | 2 | SNP | 60283 | T-C | atc-acc | new orf69 I226T, still has H300Y | PCR unpurified plaque |
| | 3 | SNP | 60282 | A-T | atc-ttc | new orf69 I226F, still has H300Y | PCR unpurified plaque |
| | 4 | SNP | 60279 | C-T | cgg-tgg | new orf69 R225W, still has E310G | PCR unpurified plaque |
| | 5 | SNP | 60280 | G-A | cgg-cag | new orf69 R225Q, still has E310G | PCR unpurified plaque |
| | 6 | SNP | 60283 | T-C | atc-acc | new orf69 I226T, still has E310G | PCR unpurified plaque |
| | 7 | SNP | 60284 | C-G | atc-atg | new orf69 I226M, still has E310G | PCR unpurified plaque |
| EcoRI-Imp2 + | 8 | SNP | 60523 | G-T | cgc-ctc | new orf69 R306L, still has G88D R90H | PCR unpurified plaque |
| EcoRI-Nlp2 dual | 9 | SNP | 60523 | G-A | cgc-cac | new orf69 R306H, still has G88D R90H | PCR unpurified plaque |
| escapers | 10 | SNP | 60515 | G-A | atg-ata | new orf69 M303I, still has G88D R90H | PCR unpurified plaque |
| | 1 | SNP | 52052 | G-C (100%, 62 count) | ggt-gct | orf59 G16A | WGS |
| | | SNP | 137542 | A-G (100%, 48 count) | gat-ggt | orf133 D109G | |
| | 2 | SNP | 60752 | G-A (91%, 65 count) | atg-ata | orf69 M382I | WGS |
| | 3 | SNP | 52057 | C-G (100%, 47 count) | cat-gat | orf59 H18D | WGS |
| | | SNP | 219699 | A-G (100%, 29 count) | atg-gtg | p61 M112V | |
| | 4 | SNP | 52052 | G-A | ggt-gat | orf59 G16D | PCR 3x plaque purified |
| | 5 | SNP | 51996 | G-T (100%, 27 count) | noncoding, -10 base from orf59 start codon | | WGS |
| | 6 | SNP | 52072 | A-C | aca-cca | orf59 T23P | PCR 3x plaque purified |
| | 7 | SNP | 52421 | G-A | cgt-cat | orf59 R139H | PCR 2x plaque purified |
| | 8 | SNP | 52108 | C-T | cgt-tgt | orf59 R35C | PCR 2x plaque purified |
| | 9 | SNP | 59824 | C-T | cca-cta | orf69 P73L | PCR 3x plaque purified |
| Tn7::Imp1, [EcoRI- | 10 | SNP | 51996 | G-A | noncoding, -10 base from orf59 start codon | | PCR 3x plaque purified |
| Imp2] | 11 | SNP | 52300 | C-A | cgt-agt | orf59 R99S | PCR 3x plaque purified |
| | 1 | SNP | 6844 | C-A (100%, 51 count) | gac-gaa | orf14 D213E | WGS |
| | | Insertion | 52123-52124 | CTG | ctg | orf59 A41 | |
| | 2 | SNP | 52033 | C-T (100%, 221 count) | ctc-ttc | orf59 L10F | WGS |
| | | SNP | 64402 | A-C (100%, 134 count) | aaa-aac | orf71 K376N | |
| | | SNP | 71928 | G-A (100%, 91 count) | gcc-gtc | orf77 A526V | |
| | | SNP | 147690 | T-G (100%, 172 count) | tcg-gcg | orf145 S139A | |
| | | SNP | 156488 | A-G (100%, 159 count) | atg-acg | orf152 M235T | |
| | 3 | SNP | 60884 | C-A (100%, 115 count) | gac-gaa | orf69 D426E | WGS |
| | 4 | SNP | 59932 | G-T (100%, 131 count) | cgt-ctt | gp69 R109L | WGS |
| | 5 | Deletion | 52124-52126 | CTG | ctg | orf59 A41 deletion | WGS |
| | | SNP | 229442 | G-A (100%, 107 count) | gcg-gca | orf232 A48A | |
| EcoRI-Imp1$_{\text{phiPA3}}$ | 6 | SNP | 52601 | G-A (100%, 63 count) | ggt-gat | orf59 G199D | WGS |
| | 1 | SNP | 52032 | C-G (100%, 315 count) | gac-gag | orf59 D9E | WGS |
| | 2 | SNP | 261894 | C-T (98%, 91 count) | acc-atc | orf287 T108I | WGS |
| | 3 | SNP | 39527 | C-A (100%, 176 count) | noncoding, -15 base from orf48 start codon | | WGS |
| | 4 | SNP | 39604 | G-T | ttg-ttt | orf48 L21F | PCR lysate |
| | 5 | SNP | 52528 | G-T (100%, 133 count) | gac-tac | orf59 D175Y | WGS |
| | 6 | SNP | 60441 | G-A (100%, 162 count) | gga-aga | orf69 G279R | WGS |
| | 7 | SNP | 39618 | C-A (100%, 156 count) | aca-aaa | orf48 T26K | WGS |
| | 8 | SNP | 39712 | A-G | ata-atg | orf48 I57M | PCR lysate |
| EcoRI-TopA | | SNP | 59830 | C-T | gct-gtt | orf69 A75V | |
